# Polyacrylamide Bead Sensors for *in vivo* Quantification of Cell-Scale Stress in Zebrafish Development

**DOI:** 10.1101/420844

**Authors:** N. Träber, K. Uhlmann, S. Girardo, G. Kesavan, K. Wagner, J. Friedrichs, R. Goswami, K. Bai, M. Brand, C. Werner, D. Balzani, J. Guck

**Affiliations:** Leibniz-Institut für Polymerforschung Dresden e. V., Hohe Str. 6, 01069 Dresden, Germany; Biotechnology Center, Center for Molecular and Cellular Bioengineering, Technische Universität Dresden, Tatzberg 47/49, 01307 Dresden, Germany; Chair of Continuum Mechanics, Ruhr-Universität Bochum, Universitätsstraße 150, 44801 Bochum, Germany; Center for Regenerative Therapies Dresden, Center for Molecular and Cellular Bioengineering, Technische Universität Dresden, Fetscherstr. 105, 01307 Dresden, Germany

## Abstract

Mechanical stress exerted and experienced by cells during tissue morphogenesis and organ formation plays an important role in embryonic development. While techniques to quantify mechanical stresses *in vitro* are available, few methods exist for studying stresses in living organisms. Here, we describe and characterize cell-like polyacrylamide (PAAm) bead sensors with well-defined elastic properties and size for *in vivo* quantification of cell-scale stresses. The beads were injected into developing zebrafish embryos and their deformations were computationally analyzed to delineate spatio-temporal local acting stresses. With this computational analysis-based cell-scale stress sensing (COMPAX) we are able to detect pulsatile pressure propagation in the developing neural rod potentially originating from polarized midline cell divisions and continuous tissue flow. COMPAX is expected to provide novel spatiotemporal insight into developmental processes at the local tissue level and to facilitate quantitative investigation and a better understanding of morphogenetic processes.

## INTRODUCTION

Mechanical signalling has been established as one of the key factors regulating cellular behaviour^1–3^and thereby driving embryonic development, morphogenesis and tissue patterning^4–11^. An important prerequisite for further progress is the availability of appropriate techniques that can either quantify stresses acting on or exerted by cells, or apply known stresses to cells to study their biological response. Many measurement techniques already exist for this purpose. Atomic force microscopy^12,13^and micropipette aspiration^14^are used as standard techniques for quantification of mechanical properties by the direct application of controlled forces to cells or tissue samples. In contrast, traction force microscopy^15^and micropillars^16,17^use deformation of the surrounding material to determine cell-generated tensional forces. Nonetheless, it is important to emphasize that these techniques enable only *in vitro* investigation of the interplay between mechanical cues or material properties and the resulting cellular behaviour. The detection of forces *in vivo* remains challenging and requires the development of appropriate tools^18–20^.

Fluorescence resonance energy transfer (FRET) sensors are increasingly used on a molecular scale to detect and quantify forces acting inside living organisms^21,22^. While this is a valuable tool to measure forces acting on individual molecules, it is unable to provide information on cell-scale stresses. With the introduction of biocompatible oil microdroplets as force transducers at the cellular level, Campas et al. (2014) have developed a pioneering method for quantifying cell-generated forces in living tissues^23^. Based on cell-induced deformations of the microdroplets, that were microinjected into the cellular space of cell aggregates and tissue explants, anisotropic normal stresses could be quantified. However, due to the incompressibility of the oil microdroplets, the estimation of isotropic tissue pressures (hydrostatic pressure) was not possible.

Recently, the concept of using calibrated spherical probes as force sensors has been refined by the introduction of elastic polyacrylamide (PAAm) microbeads to quantify compressive stress in multicellular spheroids^24^. In contrast to oil microdroplets, hydrogel probes are compressible and therefore allow quantification of local pressure changes by tracking the bead volume change. However, as the shear or elastic moduli of these PAAm beads remained undetermined, their application was limited to the identification of isotropic stresses.

In a recent study, elastic round microgels (ERMGs) made of alginate, loaded with fluorescent nanoparticles, were used to quantify isotropic and anisotropic compressive stresses in living tissues by tracking nanoparticle displacement^25^. This represents a promising method for identifying stresses, both *in vitro* and *in vivo*. The quantitative analysis of microsphere deformations rests on the definition of a stress-free spherical shape as reference configuration. Thus, the accuracy of this general approach critically depends on the availability of compressible, elastic spheres whose shape, size, and mechanical properties are well established prior to injection.

Here, we present a method that circumvents the limitations of existing cell-scale stress sensors. We validate PAAm as suitable material to create cell-like, spherical beads with a narrow size distribution using droplet microfluidics. We extensively characterize both the compressibility and elastic modulus of the beads to enable the quantification of isotropic and anisotropic stresses. This detailed determination of size and mechanical properties before injection provides the methodical basis for the *in vivo* use of the computational analysis-based cell-scale stress sensing (COMPAX). Importantly, it removes the necessity to recover the beads observed after the experiment in order to acquire knowledge of their individual stress-free reference configuration. We demonstrate the utility of COMPAX using PAAm beads by spatially and temporally quantifying for the first time the stresses acting in the developing zebrafish neural rod. We show the presence of oscillatory pressure propagation, which may potentially be generated by polarized cell divisions, and quantify compressive stress distributions within the tissue during neural rod development. COMPAX will not only provide novel spatiotemporal insight into morphogenetic processes in embryos, but can also be used for quantifying stresses in adult tissues and organoids, or when beads encounter individual cells during phagocytosis or migration.

## RESULTS

### Fabrication and mechanical characterization of PAAm beads

The production of standardized cell-like elastic PAAm beads has recently been described by our group elsewhere^26^. In order to use these beads as force sensors, we adjusted their size and elastic properties to be almost uniform and similar to those of cells. Spherical PAAm beads were prepared by controlled polymerization of acrylamide and N,N′-methylenebisacrylamide in a flow-focusing microfluidic device (left inset Fig. 1a) to obtain uniform particle size with a diameter of 17.0 µm ± 0.5 µm (mean ± SD, Fig. 1a). By altering the total monomer concentrations, the elastic modulus of the beads was adjusted to mimic the typical stiffness range of eukaryotic cells (assuming that cells can be represented as a homogeneous isotropic elastic material at the lowest order). To render the inert PAAm beads bioadhesive, and to allow a visualization of the stress response in real time, they were covalently modified with Poly-L-Lysine (PLL) conjugated with Cy3 fluorophores (PLL-Cy3) via NHS-ester, after production (right inset Fig. 1a)^26^.

Atomic force microscopy (AFM)-based colloidal probe nanoindentation was applied to determine the elastic modulus of the PAAm beads in an initial (small) strain regime and to analyze the effect of different processing steps and environmental parameters on bead stiffness. Figure 1b shows the effect of PLL-Cy3 functionalization on the elastic modulus of the PAAm beads. Unmodified PAAm beads exhibited a Young’s modulus of 1.4 ± 0.6 kPa (mean ± SD). PAAm beads which were modified with NHS-ester did not significantly increase bead elasticity (1.6 ± 0.6 kPa). The addition of PLL-Cy3 to NHS modified beads slightly increased bead elasticity to 1.8 ± 0.7 kPa. This value was then used for further analysis of the bead deformation *in vivo*. The insensitivity of the mechanical bead properties to different physiological temperatures was confirmed by AFM measurements of the same PAAm beads at temperatures from 24°C – 50°C (Fig. 1c). At all conditions tested, the elastic modulus of the beads remained constant. Young’s moduli determined using AFM were validated by numerical reconstruction of the indentation process and revealed that the reconstructed radial displacement matched the increase in bead diameter obtained by confocal microscopy during colloidal probe indentation (Supplementary Fig. 1).

Measuring the volume variation of the PLL-Cy3 PAAm beads as function of osmotic pressure using dextran solutions^24,27^was used to determine the bulk modulus (Fig. 1d). Linear fitting of the curve at small strains in the region of elastic deformation resulted in a bulk modulus of *K* = 7.7 +/- 0.2 kPa (mean ± SD). Considering the Poisson ratio *v*, which was previously determined to be 0.443^26^, the correlation between bulk modulus *K* and elastic modulus *E* can be described using the equation *E = 3_K_ (1 – 2v),* which corresponds to a Young’s modulus of 3.1 +/- 0.3 kPa. This independently confirmed the value obtained by AFM indentation. In addition, plate compression of PAAm bulk gels also verified bead elasticity at small compressions (Supplementary Fig. 2). Simultaneously, these measurements displayed a non-linear material behavior of PAAm at large strains, which precluded the application of the small strain (geometrically linear) framework. However, for moderate strains up to 15% the material response was only slightly nonlinear, which indicated the possibility to use a rather simple finite strain formulation (e.g. Neo-Hookean model) for the analysis of bead deformations.

Compressibility of the PAAm beads was further confirmed by incorporation of the beads into human mesenchymal stromal cell (MSC) aggregates (Supplementary Fig. 3). At 24 h and 48 h of culture, PLL-Cy3 functionalized PAAm beads showed significant volume reduction. After dissociation of the cells after 14 days of culture, the bead volume returned back to its initial value (shape recovery), illustrating their elastic material properties even over extended periods of time.

Together, this detailed material characterization of the PAAm beads size, compressibility and elastic modulus provide the basis for subsequent computational analysis of *in vivo* bead deformations.

**Figure 1:**
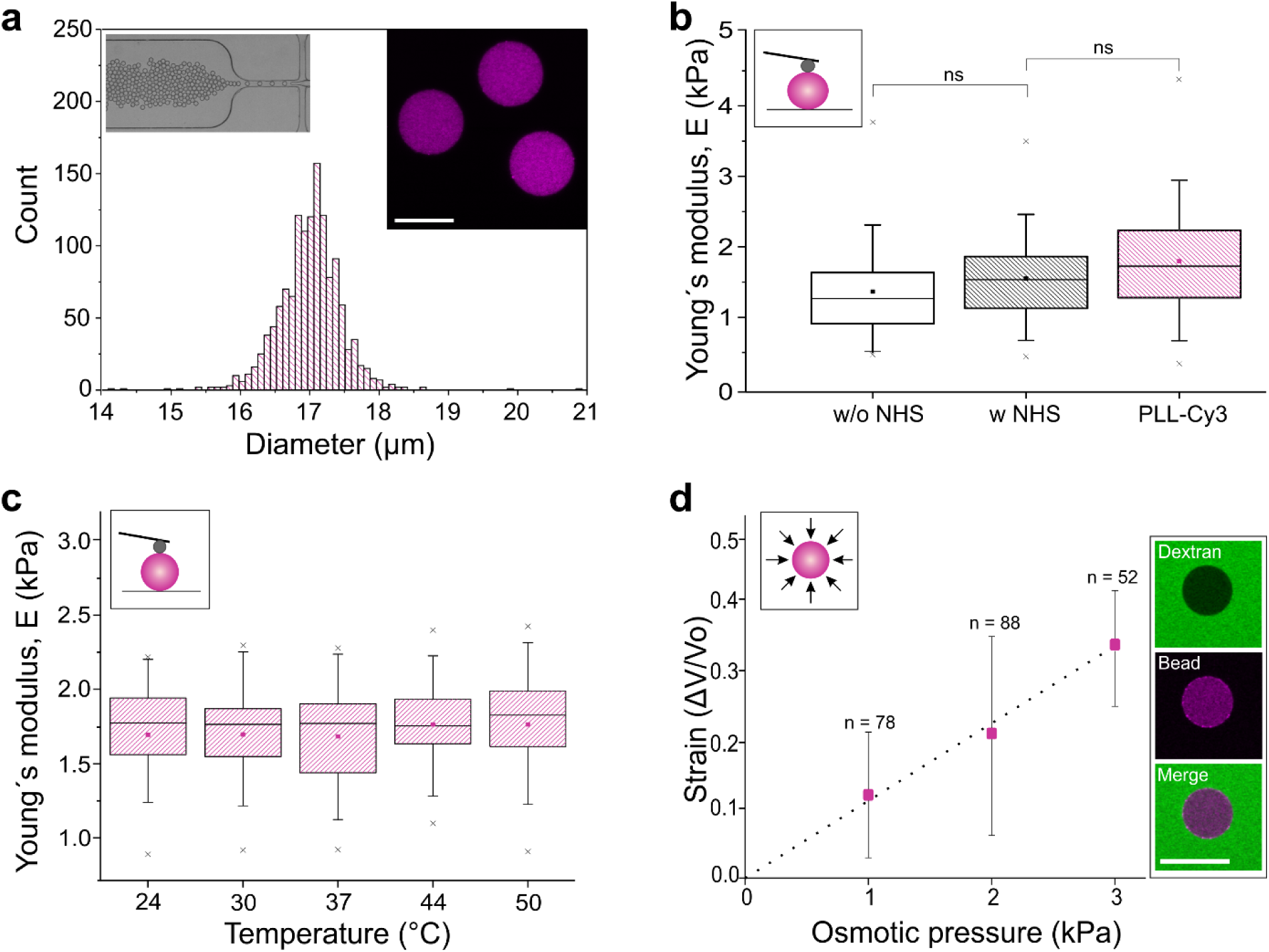
Mechanical characterization of polyacrylamide (PAAm) beads. **(a)** Size distribution of Cy3 conjugated Poly-L-lysine (PLL-Cy3) functionalized PAAm beads (*n* = 1339) determined by a macro implemented on Fiji^26^. Inset left: microscopic image of the bead production by microfluidic flow-focusing. Inset right: confocal microscopy section showing PAAm beads (magenta) functionalized with PLL-Cy3. Scale bar, 20 µm. **(b)** Young’s modulus of PAAm beads as determined by AFM-based indentation after PAAm polymerization (w/o NHS), after NHS ester modification (w NHS) and after PLL-Cy3 functionalization (*n* = 50, temperature = 24°C). **(c)** Young’s modulus stability with increasing temperature. Identical PLL-Cy3 functionalized PAAm beads (*n* = 25) were measured for each condition. **(d)** Stress-strain relation of PLL-Cy3 functionalized PAAm beads during osmotic compression. The dotted line indicates a linear fit that is used for the determination of the bulk modulus. Results are represented as mean ± SD. Inset: confocal microscopy images of a PAAm bead (magenta) in dextran solution (FITC label, green) confirming that dextran molecules (hydrodynamic radius: 27 nm) are not able to enter the polymer network (mesh size: 21 nm^26^). Scale bar, 20 µm. **(b) – (c)** The boxes are determined by the 25^th^and 75^th^percentiles. The mean is shown as filled square symbol, the median as straight line, the whiskers represent the standard deviation and the 1^st^and 99^th^percentiles are indicated by crosses. **(b) – (d)** Insets are a schematic representation of the applied methods. A homogeneous osmotic pressure compressing the bead and a spherical cantilever tip indenting the PAAm bead, respectively.

### Computational analysis-based cell-scale stress sensing

In the current study, cell-induced bead deformations are analyzed and converted to stresses using a newly developed method called computational analysis-based cell-scale stress sensing (COMPAX). COMPAX consists of three basic steps. In step one (Fig. 2a), we registered two-dimensional confocal images of the deformed PAAm beads embedded in the tissue of a zebrafish embryo. Since we were aiming to capture the detailed shape of the beads while avoiding as much noise as possible in the confocal images, the distance between the individual two-dimensional sections was chosen as 1 μm. Based on this, we constructed a three-dimensional geometry of the deformed PAAm bead and a suitable finite element (FE) discretization using the 3D visualization and processing software Avizo.

In step two (Fig. 2b) the undeformed configuration and approximate boundary conditions were estimated. Since each deformed bead can not (yet) be assigned to its undeformed configuration due to technical reasons, the undeformed state is generally not known. However, encouraged by the almost uniform fabrication of the PAAm beads resulting in narrow distributions in shape and diameter, we considered the reference geometry of the beads to be a sphere with a mean diameter of 17.0 μm. To construct the FE mesh for the respective reference configuration, we computed radial distance vectors pointing out from the surface nodes of the deformed mesh to the assumed spherical reference geometry and utilized those distance vectors as Dirichlet boundary conditions during a preprocessing FE analysis. The resulting deformed configuration was then used as the mesh for the assumed undeformed geometry of the bead.

In step three (Fig. 2c) we applied the inverse radial distance vectors as surface displacements to the mesh constructed in step 2. Then, the main FE simulation was performed to obtain the stress distribution that was assumed to mimic the one in the real PAAm bead. For this simulation we considered the Neo-Hooke material model in a geometrically nonlinear continuum mechanics setting that allowed for large displacements (see also Supplementary Fig. 1). The two material parameters *E* and *v* were estimated from the linearized situation obtained previously, i.e. they coincided with the Young’s modulus and the Poisson ratio from the linear elasticity framework that, describes the initial stress-strain response under small strains. Considering large strains, the model yielded a slightly nonlinear increase in stresses with increase in strains. This was motivated by the fact that large displacements occurred in the bead; however, the resulting stress-strain response was only slightly nonlinear in the regime of large strains up to approximately 10% (Supplementary Fig. 2). Note that strains in the beads analyzed were mostly of this level, while local maximal values reached to approximately 20%.

**Figure 2:**
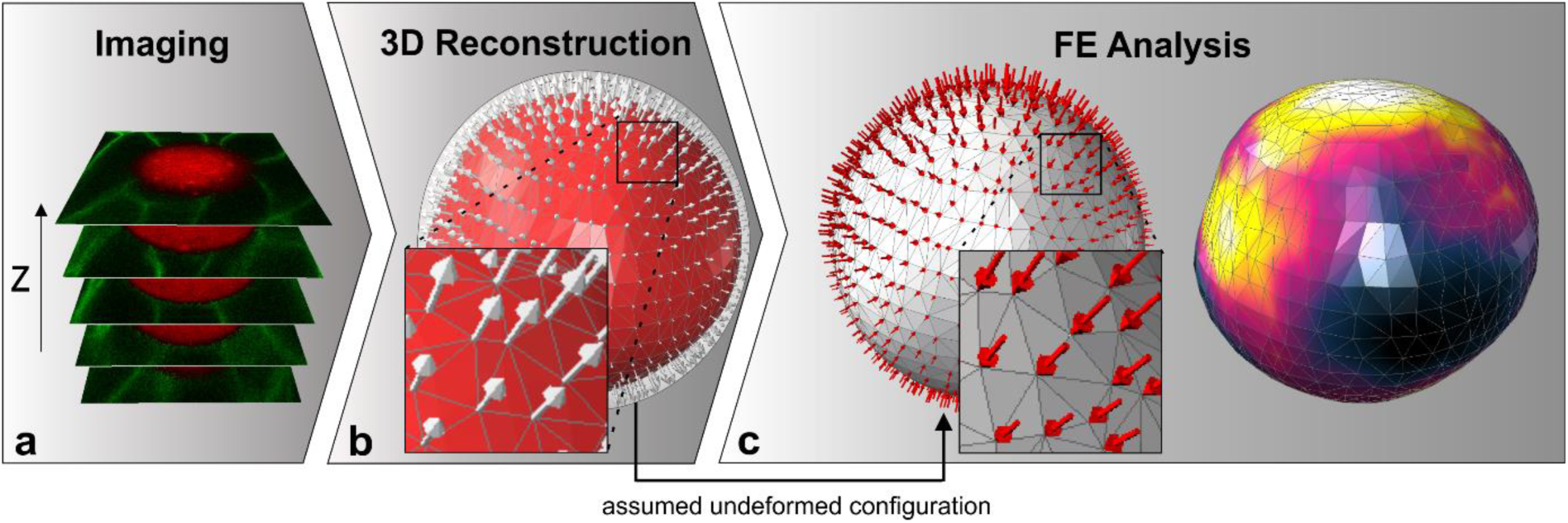
Workflow for the computational analysis of cell-scale stresses. **(a)** The deformed shape of the PAAm bead is captured by confocal microscopy. (**b**) Three-dimensional FE discretization of the undeformed configuration is constructed. For this purpose, radial distance vectors (light grey arrows) are computed pointing to the surface of the undeformed configuration assumed to be a sphere. **(c)** The inverse radial distance vectors (red arrows) are applied as surface displacement vectors in an FE analysis to compute the stresses of the deformed bead.

### Numerical validation

COMPAX constitutes a new approach to quantify cellular stresses inside soft tissues, which permits the determination of isotropic stresses as well as distinct shape changes of the beads. Therefore, a numerical validation was of mayor importance to evaluate the accuracy of the computed stress state from a qualitative and quantitative point of view. For this purpose, we analyzed several artificially created load scenarios applied to a spherical bead with a diameter of 17.0 μm, a Young’s modulus of 1.8 kPa, and a Poisson ratio of 0.443. These artificially deformed beads served as virtual experiments, that compared stresses obtained using COMPAX by analyzing the shapes resulting from the reference simulation with known data used for the simulation (Fig. 3). Note that the same diameter and material parameters were used for COMPAX as for the reference simulation such that the differences in the results should be on the order of computer accuracy in a perfect scenario.

The quality of our approach was assessed via four criteria: the local differences in surface pressure σ_Pres_, the magnitude and direction of the volumetric mean of the principal stresses 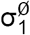, 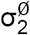 and 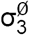, and the volumetric mean of pressure 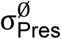. Here, pressure refers to negative hydrostatic stress, defined as the arithmetic mean of normal stresses. Positive values of the pressure indicate compressive isotropic stresses. As a plausibility check, we first compared the results for a homogeneous surface pressure of 1000 Pa, where the resulting displacements are indeed radial. The resulting stresses of COMPAX differed from the virtual data by values in the order of computer accuracy, implying that the method had been correctly implemented. In a first relevant test scenario, we defined a periodic surface pressure whose intensity varied sinusoidally with circle coordinates angle φ in the x-y-plane from a minimum of 600 Pa to a maximum of 1000 Pa with a wavelength of 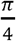. In this case, the comparison revealed an excellent agreement between virtual data and the COMPAX-method (Fig. 3a). The second test scenario considered a quadratic distribution of surface pressure along one rotation axis of the bead with a maximum pressure of 1000 Pa at the equator and a minimum pressure of 600 Pa at the poles. The contour plots exhibited some local quantitative differences, while also the values of the volumetric means of the principal stresses varied (Fig. 3b). Nevertheless, the direction of principal stresses as well as the mean pressure showed good conformity. In a final load scenario, we combined a homogeneous surface pressure of 800 Pa with two different shear loads of 120 Pa. Each of the two loads was applied tangentially on one of the two halves of the bead. The first and second loads were oriented from the north to the south pole and from the east to the west pole, respectively. Comparing the contour plot in Fig. 3c, significant quantitative and qualitative differences in the local pressure distributions could be observed, even though the directions of the principal stresses and volumetric mean of the pressure remained comparable.

These artificial situations did not yet include potential errors resulting from measurement uncertainties e.g., associated with the shape of the undeformed bead and the material parameters. Therefore, an uncertainty analysis was performed for diameter and Young’s modulus, which showed an influence of up to 30% standard deviation in volumetric mean pressure as a result of measured differences in these two parameters (Supplementary Fig. 4). Taken together, COMPAX was less reliable when extracting surface shear and larger non-radial deformations, which led to local deviations in stress computation and simultaneously affected the values of principal stresses. However, we obtained very satisfying results for the determination of mean pressure and directions of the principle stresses, which were thus primarily analyzed in the experiments.

**Figure 3:**
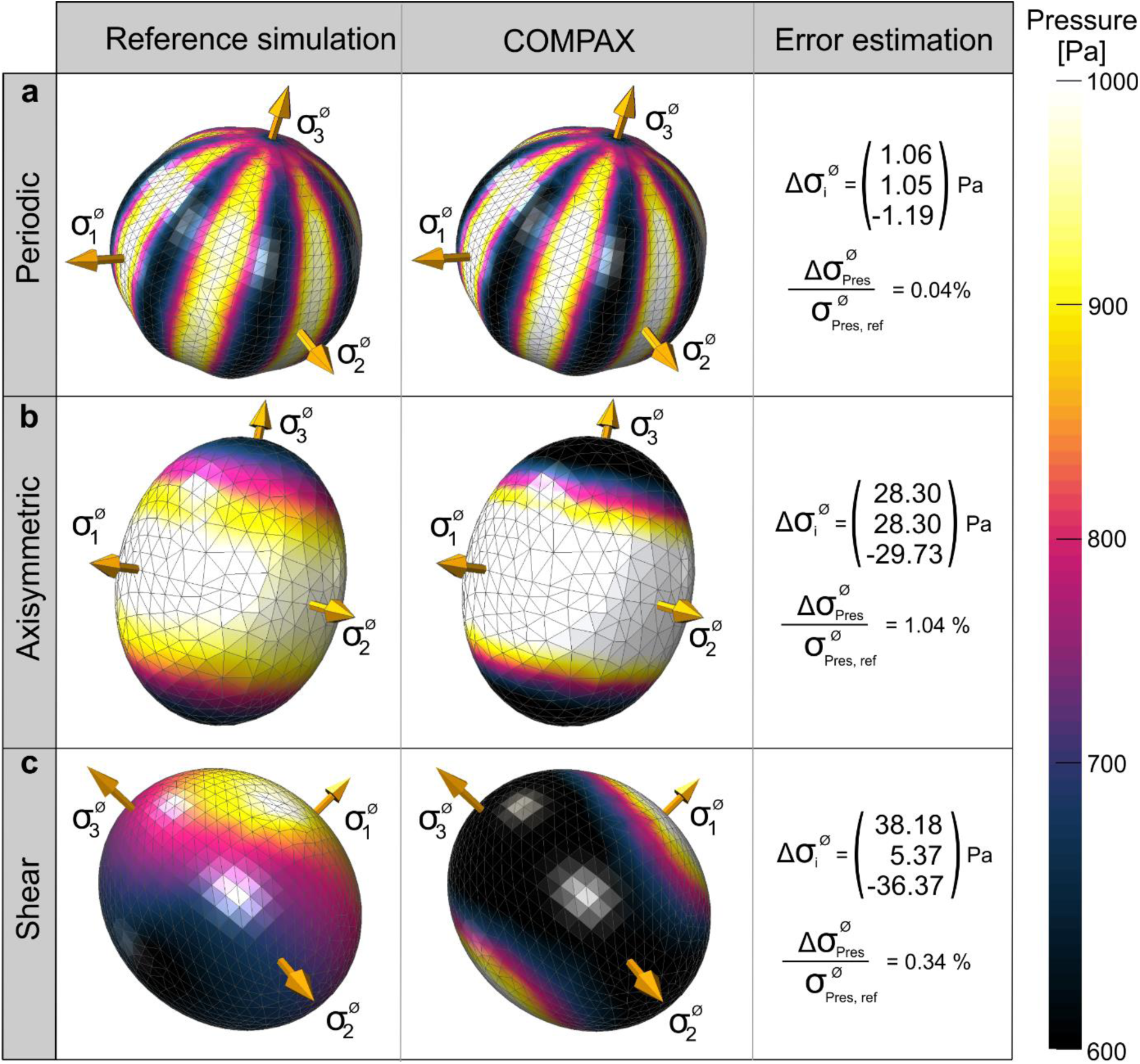
Method validation using analysis of numerically deformed beads. 3D contour plots of pressure and principal stresses in the reference simulation (left) and results of the COMPAX method (middle). Reference values based on the volumetric mean of the Cauchy stress tensor (right). **(a)** The application of periodic surface pressure shows nearly same results for both computations. **(b)** Larger non-radial displacements lead to local deviations in the contour plot. **(c)** Stresses reveal inverse local values for the numerical simulation of surface shear, but nearly no difference in mean pressure.

### Quantifying cell scale stresses during zebrafish development

Cell-generated forces that emerge during neural rod formation in zebrafish embryos were quantified by computationally reconstructing the deformations of embedded PLL-Cy3 PAAm beads. The beads were microinjected into the developing neural plate at the tailbud stage (10 hours post fertilization (hpf)) and time-lapse confocal microscopy was used to visualize three-dimensional shape changes of the beads in the neuro-epithelium (Fig. 4).

**Figure 4:**
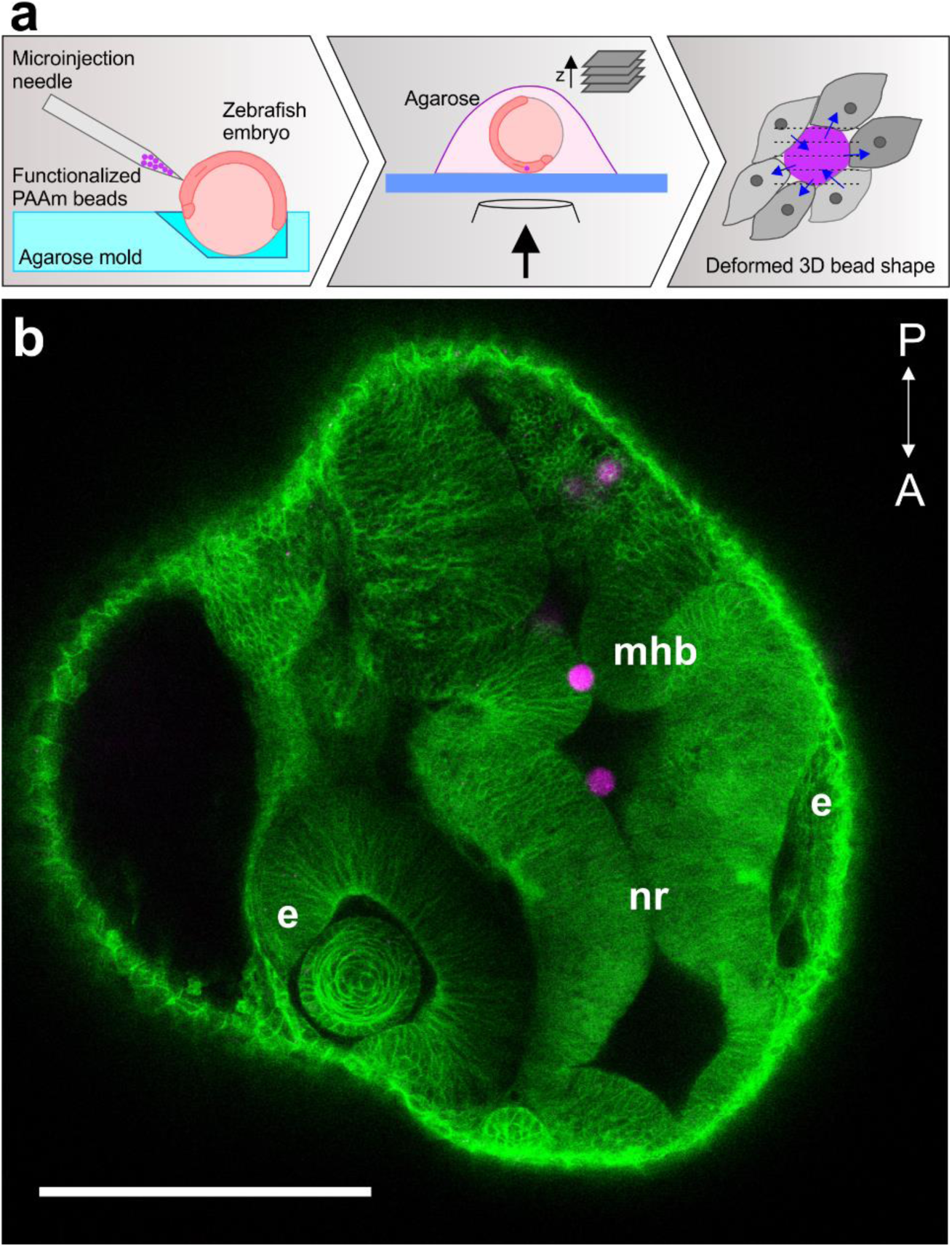
PAAm bead microinjection into zebrafish embryos. **(a)** Schematic illustration of the microinjection and imaging procedures. The embryo was immobilized in an agarose mold to facilitate upward orientation of the animal pole for microinjection. PLL-Cy3 PAAm beads were injected at the bud stage of zebrafish development (10 hpf). For confocal imaging of cell-force-induced bead deformations, the embryo was embedded in 1% low melting agarose. **(b)** PAAm bead trapped between the left and right side of a uniformly opened neural rod (nr) in the region of the midbrain-hindbrain boundary (mhb) of a zebrafish embryo at prim-15 stage (30 hpf). Developing eyes (e) and structures of the central nervous system are recognizable. Anterior (A) - posterior (P) direction is marked in the image. Scale bar, 200 µm; the PLL-Cy3 PAAm beads are magenta colored and the cell membranes are GFP-labeled (green).

First, we analyzed cell-induced deformations of a bead embedded in the developing neural rod of a zebrafish embryo. Bead and embryo were imaged for different periods of time at 14 hpf, 15 hpf, and 20 hpf (Fig. 5a). Throughout the imaging period, the bead remained constantly positioned in the basal region of neural rod adjacent to the developing otic placodes of the embryo. COMPAX at the indicated time points showed that the surrounding cells exerted forces on the bead that induced its deformation (Fig. 5b). Here, we wanted to distinguish between local stresses on distinct areas of a bead and the global volumetric mean of pressure. While local tensile isotropic stresses (negative pressure values) were initially detected at 14 hpf, volumetric compression (positive pressure values) dominated at later time points. The temporal evolution of volumetric mean of the pressure for each time period, as well as the corresponding volume of the PAAm bead, is shown in Supplementary Fig. 5a. Additionally, COMPAX-based rendering of local compressive stresses can be related to the activity of adjacent cells, where cell spreading/division caused greater local compressive deformations (Fig. 5c). To quantify prevailing stress distributions during neural rod formation, we computed the normal compressive stresses acting in the x-y-plane as a function of the angle φ (Fig. 5d). For better visualization, the normal compressive stresses were normalized to the maximal normal compressive stress value. Also, the differences to the mean normal stress, computed for the corresponding current configuration, were amplified by a scaling factor of five. We observed a continuous pulsatile pressure propagation pattern in the basal part of the neural rod during the evaluated periods. The intensity of pressure oscillations decreased with time but were clearly detectable for the entire period of evaluation. Decreased changes between stress distributions of serial time points implied the dominance of the global stress state over stress changes caused by local developments. Furthermore, deformation of the beads indicated a change in stress orientation during the first imaging period at 14 hpf, as evidenced by the change in the direction of the maximum compressive normal stress amplitude. Stress orientation remained unaltered during subsequent periods.

**Figure 5:**
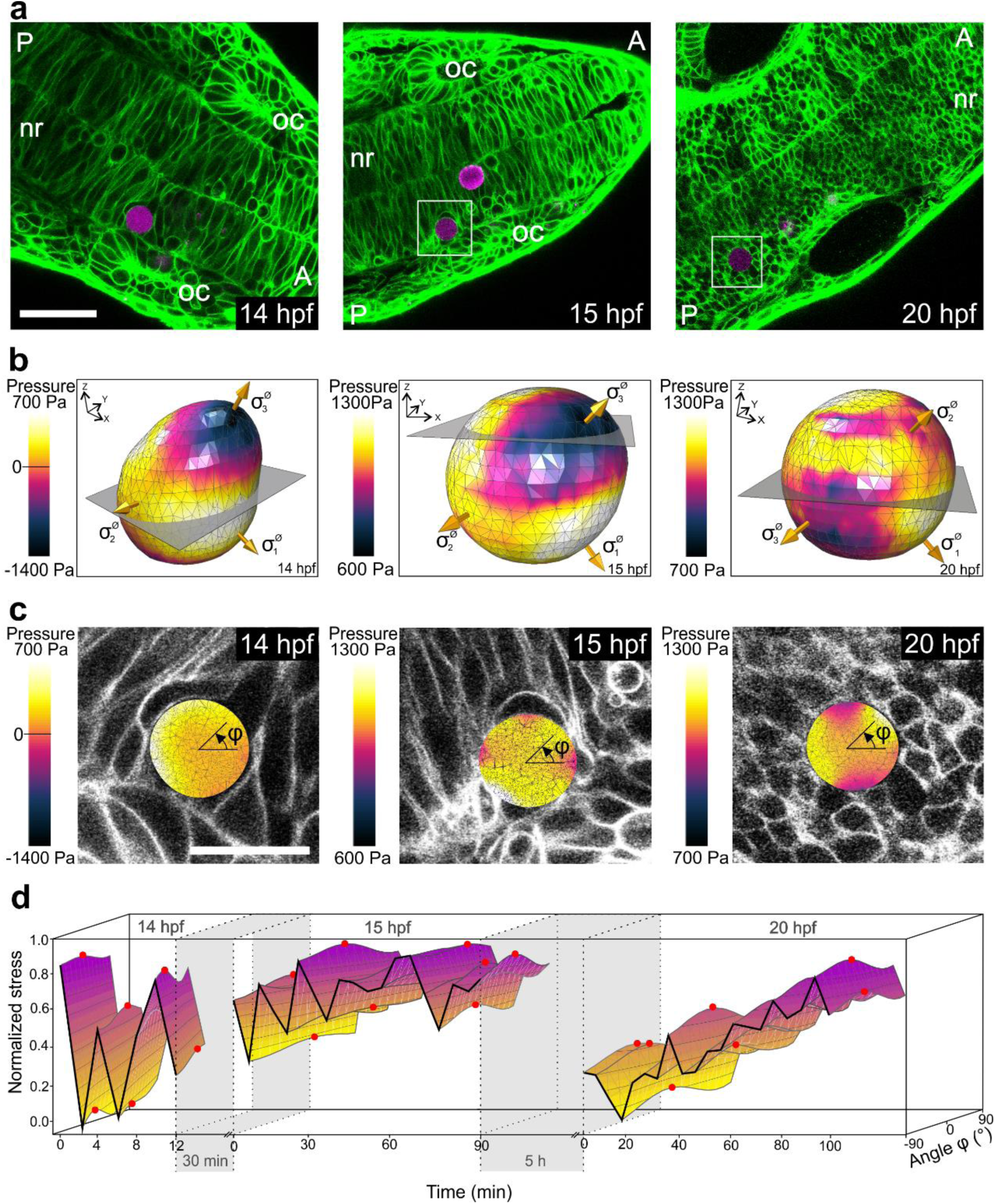
In vivo quantification of cellular compressive stresses during zebrafish neural rod formation. **(a)** Confocal sections of the developing neural rod (nr) of a zebrafish embryo after bead injection at 14 hpf, 15 hpf, and 20 hpf. The PLL-Cy3 PAAm bead (magenta label; white box) is embedded between neural progenitor cells at the basal part of the neural rod, reframed by the developing otic placodes (oc). Anterior (A) and posterior (P) direction is always marked in the image. The plasma membrane of all cells in the zebrafish embryo is marked in green. Scale bar, 50 µm. **(b)** Representative contour plot of pressure on the PAAm bead at the beginning of the imaging period at 14 hpf, 15 hpf, and 20 hpf. The golden arrows indicate the directions of the principal stresses. The grey slices represent the positions of the confocal planes depicted in (c). **(c)** Overlay of the cross-section of the contour plot (presented in b) and the corresponding confocal plane of the z-stack at 14 hpf, 15 hpf, and 20 hpf. The angle *φ* represents the orientation of the normal compressive stress depicted in d. **(d)** 3D representation of the normalized amplified compressive stress distribution for the entire imaging period at 14 hpf, 15 hpf, and 20 hpf. Maximum amplitudes of normal compressive stresses are displayed as red dots. In this and all subsequent figures, the time periods between the measurements are indicated by grey areas.

This occurrence of periodic pressure fluctuations during zebrafish development was confirmed by microinjecting additional beads into the neural plate and tracking their deformations at identical developmental stages (biological replicates). A PAAm bead at a position comparable to the bead shown in Figure 5 — in the developing structure of the otic placode (basally localized) — also exhibited oscillatory compressive normal stresses at 14 hpf and 15 hpf (Fig. 6a, Supplementary Fig. 5b). In contrast, a bead positioned close to the midline (apically localized) of the neural rod within the same embryo (Fig 6b, Supplementary Fig. 5c) was simultaneously exposed to pulsatile tensile stresses. Deformation patterns of another bead embedded near the midline in another embryo (Fig. 6c, Supplementary Fig. 5d) also confirmed the overall trend of oscillatory stress propagation, but showed compressive stresses only at 14 hpf, which might be explained by the continuous movement of this bead from the midline towards the basal border of the neural rod (inset Fig. 6c, Supplementary Fig. 5e). To visualize prevailing forces after the opening of ventricles in the neural rod, deformations of a bead located at the ventricle at 19 hpf were imaged (Fig. 6d). The bead was surrounded by cells on two sides but unattached on the other two sides. COMPAX analysis revealed oscillating compressive stresses within the developing cerebral rod. Overall, these results suggest the presence of spatially varying oscillatory stresses during development of the zebrafish neural rod.

**Figure 6:**
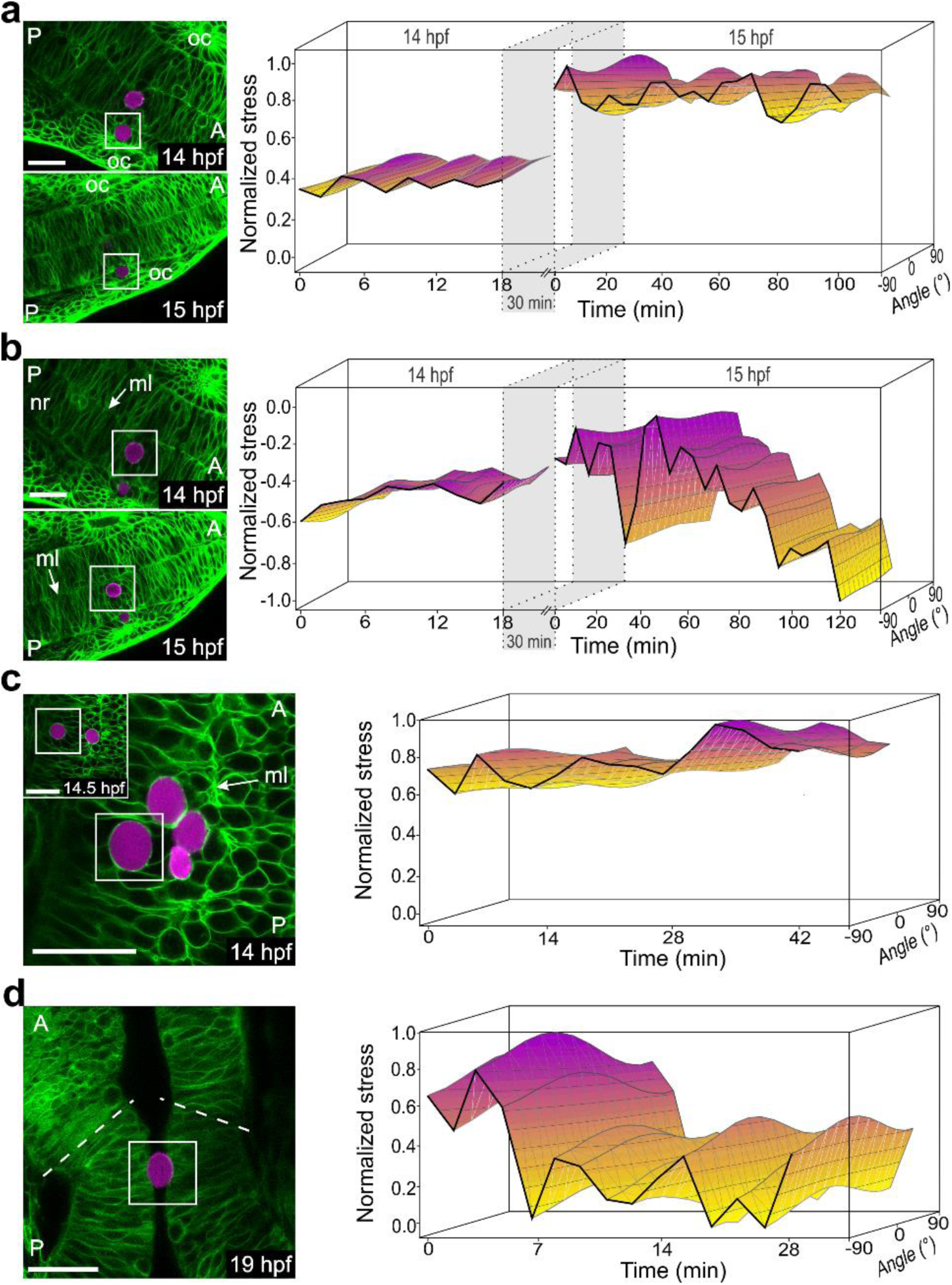
Spatial and temporal normal compressive stress variations within the zebrafish neural rod during development. **(a)** Left panel: Confocal section of a PLL-Cy3 PAAm bead (white box) within the neural rod embedded between cells of the otic placode (oc) at the 10-somite stage (14 hpf) and the 12-somite stage (15 hpf). Right panel: 3D representation of the normalized amplified compressive stress distribution for the entire imaging period at 14 hpf and 15 hpf, respectively. **(b)** Left panel: Confocal section of a PLL-Cy3 PAAm bead (white box) embedded (in the same embryo as shown in panel a) close to the midline (ml) of the neural rod (nr) at the 10-somite stage and the 12-somite stage. Right panel: 3D representation of the normalized amplified compressive stress distribution over the entire imaging period at 14 hpf and 15 hpf, respectively. Note that negative normalized stress values correspond with tensile stresses. **(c)** Left panel: Confocal section of a PLL-Cy3 PAAm bead (white box) close to the midline (ml) of the neural rod at 14 hpf. Inset: Identical bead after movement towards the basal part of the neural rod at the end of the imaging period at 14.5 hpf. Right panel: 3D representation of the normalized amplified compressive stress distribution for the entire imaging period. **(d)** Left panel: Confocal section of a PLL-Cy3 PAAm bead trapped in the developing midbrain-hindrain (mbh) of the neural rod at 19 hpf. The dashed lines indicate the border of the mbh. Right panel: 3D representation of the normalized amplified compressive stress distribution for the entire imaging period. **(a) - (d)** Confocal images: anterior (A) and posterior (P) direction is always marked in the image. PLL-Cy3 PAAm bead (magenta label) and cell membranes (GFP labeled; green). Scale bar, 30 µm.

## DISCUSSION

The present work has demonstrated that PLL-Cy3 functionalized PAAm beads can be efficiently injected into living tissue/embryos to quantitatively determine *in vivo* cellular stresses. By numerically analyzing bead deformations that deviate from the spherical shape of the reference configuration, data on pressure changes as well as the direction of main shape changes can be extracted for the first time. Detailed mechanical characterization of the beads and elaborate computational analysis of *in vivo* bead deformation revealed pulsatile stresses during zebrafish embryo development.

The preparation of PAAm beads using standardized microfluidic flow focusing^26^resulted in an extremely narrow size distribution which, to the best of our knowledge, has not been achieved in other studies. In general, computation of bead shape changes is based on the knowledge of the stress-free state as reference. Mohagheghian et al.^25^have shown that this reference configuration could be accurately determined by cell removal during *in vitro* application of alginate microspheres. However, as this was not realizable *in vivo*, the initial state after injection was used to define the stress-free state, which introduced uncertainties in stress calculation. In contrast, the size uniformity of the PAAm beads used here was essential to render COMPAX insensitive to variations in initial size and permitted the application of this approach to living developing tissues/animals over extended periods of time.

The elastic modulus of the PAAm beads (∼ 2 kPa) was similar to that of cells or tissues which, once injected into a living tissue/embryo, allowed the surrounding cells to deform the beads, and thus enabled sensitive cell force studies at different stages of zebrafish embryonic development. Since knowledge of elastic bead properties is an essential prerequisite for a reliable stress analysis, we performed extensive material characterization measurements directly on the PAAm beads. AFM-based colloidal probe indentation was used to determine the Young’s modulus of the beads at various stages of the PLL-Cy3 functionalization process. The average Young’s modulus of NHS-modified beads slightly increased after PLL-Cy3 modification due to electrostatic interactions (additional crosslinking points). The PLL-Cy3 enhances interactions with the cell membranes (negatively charged ions) and its fluorescent label enables the detection of the beads with confocal microscopy. Modification of the gel with PLL introduced positive charges that may have altered bead elasticity by introducing additional crosslinking points into the polymer network^28^. Once the beads were modified with PLL-Cy3, the whole beads were homogenously fluorescent and displayed stable stiffness at temperatures ranging from room temperature to 50°C.

In a majority of studies on the use of colloidal microgels as stress sensors, mechanical characterization of the test system was performed on bulk gels^25,29^. Here, we have directly compared bead properties at the macro- and microscale. Pressure tests on PAAm bulk gels (identical composition as the beads) were performed and revealed similar initial elastic moduli as determined by AFM-based indentation. Nevertheless, bulk gel tests exhibited minor dispersions in curve shape compared to AFM measurements, which could be caused by variations in sensitivity to the porosity of the gels during macro- and microscale responses. Moreover, a comparison of axial material behavior of PAAm gels with the Neo-Hookean material law showed conformity for strains up to 10%. Deformations of the PAAm beads observed were within that range, which justifies the application of the Neo-Hookean material law. However, a new non-linear elastic material law needs to be developed to capture with equal accuracy larger deformations that were observed locally in the beads. This conclusion is contrary to the conventional approach of previous studies, which describe hydrogel stress sensors as linear elastic materials. The resulting numerical simulations were restricted to being geometrically linear — a scenario that did not allow for an appropriate incorporation of large strains. Additionally, bulk tests have demonstrated that further investigations of the material at the microscale could facilitate development of a more detailed understanding of mechanical behavior. As AFM-based colloidal probe indentation only analyzed the Young’s modulus for deformations less than 10%, the rising slope of the strain-stress curve from the bulk tests could not be captured. Both, macroscopic and microscopic material characterization were requisite for a detailed understanding of the material behavior which enabled us to constrain uncertainties during computations of isotropic and anisotropic stresses to a minimum. To the author’s knowledge, comparable studies have not yet achieved a similar accuracy during quantification of intracellular stresses.

Numerical validation of COMPAX exhibited good results for all load scenarios with four different criteria, along with significant accuracy during determination of principal stress orientation and the mean pressure. Only shear stress distribution could not be captured quantitatively using COMPAX, which can be explained by discrepancies of the real displacement direction from the assumed radial orientation. To increase the accuracy of the method, fluorescent and highly dispersed markers could be embedded within the beads to enable a point-wise reconstruction of displacement vectors, at least at the marker positions, and to obtain volume displacement information in addition to surface displacements. A similar approach for reconstructing shape deformations using fluorescent markers has already been used for the ERMG method^25^. However, because of the lack of a reference configuration corresponding to the shape of the stress-free bead, the marker could not be utilized to obtain detailed information on displacements. Further, another approach will be required to map the markers in the deformed configuration to corresponding markers in the undeformed configuration. For this purpose, a few different markers with a predefined distance between each other in order to re-identify the orientation of the bead in the deformed state and to map the highly dispersed markers in the undeformed configuration. This method modification could serve to identify the exact reference configuration of the undeformed initial bead and would constitute a substantial improvement. Nonetheless, since the average bead diameter is used to construct an undeformed sphere, which was assumed to be the reference geometry for the main FE analysis, uncertainty in the bead diameter had to be considered. The narrow size distribution of the fabricated PAAm beads enabled us to constrain the standard deviation of the related uncertainty analysis to 30%. The implantation of fluorescent markers could reduce this variance even further. Initial observations on tissue stress distributions in zebrafish embryos were made possible by ERMGs made of alginate injected into the blastula and early gastrulation stages of embryonic development^25^. In that study, the authors provided evidence for spatial differences in tensile and compressive stresses during the early stages of tissue development. As an extension of the previous study, we now show that PAAm beads can be used to quantify changes in stresses generated by cells during zebrafish neurulation, a dynamic morphogenetic process that transforms the neural plate into the neural rod. We also show that tensile and compressive stresses occur simultaneously at different positions within the developing neural rod and that they coincide with observations derived from early developmental studies that used ERMGs. Importantly, both studies indicate that spatial variations can arise in prevailing stresses during tissue formation. In the present study, we also provide evidence that compressive stresses might dominate neural rod formation between 14 hpf and 20 hpf. Such compressive stresses are attributable to persistent and dense tissue packing and associated feedback control of neurogenesis at the apical-basal polarity axis^30^.

Using PAAm bead stress sensors we show, for the first time, differences in spatial and temporal stress distributions not only during gastrulation, but also directly within the tissue during zebrafish neural rod formation. The observed pulsatile variations in relative stress fields during neurulation may result from polarized cell divisions, especially for beads located in the neural rod at 14 hpf and 15 hpf. Such oriented cell divisions occur during the neural keel-rod period of zebrafish neurulation and induce a uniform distribution of polarity in the neural rod through synchronized cell divisions at the midline^31–33^. As neurulation progresses, the number of cells gradually increases and, when combined with general cell shape changes and the curvature of the neural plate, can explain the observed oscillatory stress distribution. Furthermore, our observations also imply that the influence of single cell divisions related to the entire development of the fish probably decline with time as we show dominance of global stress state over stress changes due to local developments and only minor changes in stress distributions during serial time points.

In conclusion, we have used PAAm beads and COMPAX to quantify compressional forces in real-time and estimate the direction of main shape changes during neural rod formation *in vivo* in zebrafish embryos. The same principle can be applied to other fields of application in order to investigate prevailing stresses *in vivo* and *in vitro*^2,34^. From a developmental biology perspective, further investigation of stresses acting in tissues/organisms during early or later development as well in adult animals can be addressed using this system. Moreover, exploring stress changes in cultured cells, organoids, or monolayers will provide further knowledge on mechanical aspects of cell-cell interactions. Also at a single cell level, processes such as migration or phagocytosis, can be detected when the beads encounter individual cells. The versatility of this approach has been recently exemplified by Vorselen et al., who have evaluated the influence of target rigidity on phagocytosis and associated force generation by macrophages. We predict that, many other applications can be envisioned where the quantification of cell-scale stresses will improve our understanding of the role of mechanics in biology and medicine.

## METHODS

### Fabrication and modification of PAAm beads

The fabrication of PAAm beads using a microfluidic device as well as their subsequent functionalization with PLL has been described elsewhere in detail by Girado et al^26^. Briefly, the generation of PAAm beads, which were sufficiently compliant to allow cell force induced deformations and enabled a convenient handling during microinjection at the same time, was achieved by using a total monomer concentration of acrylamide (AAm) and bis-acrylamide (BIS) of 7.9%. During the production process PAAm droplets were functionalized with NHS-ester to enable their modification with Poly-L-Lysine (PLL) conjugated with Cy3 fluorophores (NANOCS). A PLL-Cy3 concentration of 28 pg/bead was used to enable a homogeneous protein functionalization.

### Atomic Force Microscopy (AFM)

The AFM indentation measurements were performed using a Nanowizard I AFM (JPK Instruments) mounted on an inverted optical microscope (Axiovert 200, Zeiss). A tipless cantilever (Arrow TL-1, nominal spring constant k = 0.035 - 0.045 N/m, NanoAndMore GmbH), equipped with a polystyrene microsphere (diameter: 5 µm, Microparticles GmbH), was used. For gluing the microsphere to the end of the cantilever, a two-component epoxy glue (Araldite) was used. The cantilever was calibrated by the thermal noise method before the experiment. For indentation, the cantilever tip was aligned over the center of the bead and individual force-distance curves were acquired with an approach velocity of 5 µm/s and with a contact force of 2 nN (typical indentation depth: 1 µm). Extracting the Young’s modulus of the beads was realized by fitting the approach force-distance curve with the Hertz model for an spherical indenter and applying a double-contact correction considering an additional deformation derived from the counter pressure at the bottom side of the bead during indentation^26,35,36^. A cell-adhesive protein solution (CellTak, Cell and Tissue Adhesive, Corning) was used to immobilize the PAAm beads on the bottom of a petri dish to prevent bead motion during the experiment. The Young’s modulus was determined using the JPK data processing software (JPK Instruments). The measurements were executed in PBS and at room temperature, except the temperature stability tests where the temperature was set to 24°C, 30°C, 37°C, 44°C and 50°C, respectively. For AFM measurements combined with confocal fluorescent microscopy the identical AFM set up and parameters were used in combination with a confocal microscope (Zeiss 510 Meta).

### Bulk modulus measurements of PAAm beads

Compressive osmotic stress for the determination of the bulk modulus of the PAAm beads was induced by PBS supplemented with fluorescein labeled dextran (molecular weight of 70 kDa, Sigma Aldrich 52471-1G). Well-defined amounts of dextran were utilized to ensure a controlled application of osmotic stress^24,27^. The three-dimensional shape of the PAAm beads was captured using an inverted confocal microscope (Zeiss LSM700) before and 30 min after exposing to the dextran solution, and the volume of the PAAm beads was analyzed using the open source software Fiji (3D image counter)^37^.

### Rheometer

For rheological measurements a plate rheometer (ARES Rheometer, Rhemetric Scientific) was used. Fully swollen PAAm bulk gels (monomer concentration: 7.9%, gel diameter of 1 cm) were compressed until 60% strain at room temperature.

### MSC aggregate formation

Human telomerase reverse transcriptase (hTERT) immortalized mesenchymal stromal cells (MSCs) were transduced according to Girardo et al^26^. and maintained under humidified 5% CO_2_ atmosphere in low-glucose Dulbecco’s modified Eagle medium (DMEM, Gibco) enriched with 10% fetal bovine serum (Gibco, Invitrogen). For multicellular aggregate formation, MSCs in suspension were mixed with PLL-Cy3 functionalized PAAm beads and cultured in form of 70 µl droplets on inverted petri dish lids for 24 h, 48 h and 14 days according to the classical hanging drop method.

### Numerical Simulations

Confocal images were imported to the 3D visualization and processing software Avizo. Therein, the bead was segmented in every image utilizing segmentation tools e. g., thresholding. Based on the segmentation the 3D geometry was interpolated and, subsequently, the FE mesh was constructed. All simulations were conducted in ABAQUS, a commercial finite element software, which provides the Neo-Hookean material law as internal material subroutine.

All stress computations were based on the Cauchy stress tensor σ. Contour plots of PAAm microbeads show the distribution of surface pressure, which is defined as 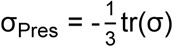. Further evaluations of the COMPAX-method were based on the volumetric mean of the Cauchy stress tensor σ, which was computed by

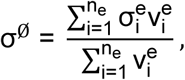

where σ^e^is the Cauchy stress tensor at a single integration point within the finite elements and v^e^is the corresponding volume. We label the principal stress of the volumetric mean of the Cauchy stress tensor to be σ_i_^e^ with i = [1,3] and the volumetric mean of the pressure 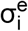.

The values of comparison 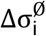 and 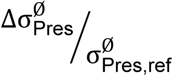 which were involved in the numerical validation, are defined as

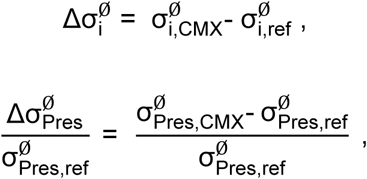

where (·)_ref_ and (·)_CMX_ constitute the labeling of values related to the reference simulation and the COMPAX-method, respectively.

### Zebrafish strain and maintenance

Zebrafish (Danio rerio) embryos and adults were obtained, raised, and maintained as described previously^38,39^. Embryos were staged as hours post fertilization (hpf)^40^. To obtain membrane-tagged eGFP embryos, adults from the transgenic strain Tg(Bactin:HRas-EGFP) (ZDB-ALT-061107-2)^41^were outcrossed with those from wild type AB strain. Furthermore, a transgenic line which expressed a nucleus-targeted venus fluorescent protein at the otx2 locus (knock-in) was established using the CRISPR knock-in strategy as previously described^42^; this line was used to identify the midbrain hindbrain boundary.

All animal experiments were carried out in accordance with animal welfare laws and local authority requirements (Landesdirektion Sachsen). Protocols for the generation (24-9168.11-1/2013-14) and maintenance of (DD24-5131/346/11 and DD24-5131/346/12), and experimentation with transgenic animals (24-9168.24-1/2014-4) were also appropriately approved (Landesdirektion Sachsen, Germany).

### Zebrafish microinjection

The embryos were grown in a 28.5 °C incubator, dechorionated at 10 hpf using pronase (Sigma-Aldrich, P8811) and incubated in calcium-free ringer solution (recipe from the zebrafish information network (ZFIN) database; http://zfin.org) for 30 minutes. The bead implantation was carried out in the same solution. Standard borosilicate capillaries without filament (World Precision Instruments) were pulled using a micropipette puller (Sutter Instruments). The tips were cut to obtain an opening of about 15 µm and back-loaded with the injection solution containing hydrogel beads. The beads were implanted into the developing neural plate close to the prospective midbrain hindbrain boundary using a microinjector system (PV820, pneumatic picopump, World Precision Instruments). The injected embryos were incubated in E2 buffer (recipe from ZFIN protocol) until imaging. The implantation of the beads did not result in any morphological or developmental abnormalities. Fish were pre-screened for successful implantation in the desired location. The selected embryos were mounted as described below and imaged.

### Embryo mounting and imaging

Embryos with beads in the desired location were anesthetized using MS-222 (Sigma Aldrich, A5040) in E2 solution and dorsally mounted on a glass-bottomed petridish (MatTek) in 1% low melting agarose prepared in E2 solution (Fig. 4a). Time-lapse images were acquired using an inverted confocal microscope (Zeiss LSM780) using a 40x water immersion objective (NA 1.2) with laser lines 488 nm for eGFP and 561 nm for Cy3. Images were analyzed using the open source software FIJI^37^.

### Statistical analysis

The bin widths of the histogram (Fig. 1a) was chosen according to the Freedman-Diaconis rule^43^. In the box plots (Fig. 1b, c; Supplementary Fig. 4b) the mean is shown as filled square symbol, the median as straight line, and the boxes are determined by the 25^th^and 75^th^percentiles. The whiskers represent the standard deviation and the 1^st^and 99^th^percentiles are indicated by crosses in Fig. 1b and c. In Fig. 1d data points with error bars indicate mean ± standard deviation. The number of measurements (*n*) is given at the respective data point boxes (Fig. 1d; Supplementary Fig. 4b) and in the figure description (Fig. 1b, c), respectively. The data in Fig. 1b and Supplementary Fig. 4b was analyzed using OriginPro Software. Both data sets rejected normally (Shapiro-Wilk test). To evaluate the statistical difference, the non-parametric Wilcoxon-Mann-Whitney test was used. The asterisks define the statistical difference as follows: *** *P* < 0.001.

## ACHKNOWLEDGEMENTS

We thank the Microstructure Facility and the Light Microscopy Facility (partly funded by the State of Saxony and the European Fund for Regional Development – EFRE) of the Center for Molecular and Cellular Bioengineering of the TU Dresden for bead production and for technical support with confocal microscopy, respectively. We thank Angela Jacobi (Biotechnology Center, TUD) for providing human telomerase reverse transcriptase (hTERT) immortalized mesenchymal stromal cells (MSCs). We thank Petra Welzel (IPF, Dresden) for help with the rheometer. We thank Elisabeth Fischer-Friedrich (Biotechnology Center, TUD) for letting us use the AFM combined with a confocal microscope. We thank Marika Fischer, Jitka Michling and Daniela Mögel for dedicated zebrafish care. We thank Dr. Vasuprada Iyengar for language and content editing. This work was also supported by an ERC advanced grant (Zf-BrainReg) and project grants of the German Research Foundation (Deutsche Forschungsgemeinschaft, project number BR 1746/6-1 and BR 1746/3) to M.B. We gratefully acknowledge financial support from the Alexander von Humboldt Foundation (Alexander von Humboldt Professorship to J.G.)

## AUTHOR CONTRIBUTION

J.G., D.B., M. B., and C.W. supervised and defined the project; N.T., G.K., and K.B. designed the bead injection routine; N.T. microinjected the beads in zebrafish embryos; K.U. and D.B. developed the computational analysis, N.T. and K.W. mechanically characterized the beads; N.T. and J.F. performed MSC aggregate experiments; S.G. designed and R.G. prepared the PAAm beads; G.K. and M.B. provided zebrafish embryos and helped with data interpretation; N.T. and K.U. wrote the paper with the help of D.B. and J.G.

